# Deep Normative Tractometry for Identifying Joint White Matter Macro- and Micro-structural Abnormalities in Alzheimer’s Disease

**DOI:** 10.1101/2024.02.05.578943

**Authors:** Yixue Feng, Bramsh Q. Chandio, Julio E. Villalon-Reina, Sebastian Benavidez, Tamoghna Chattopadhyay, Sasha Chehrzadeh, Emily Laltoo, Sophia I. Thomopoulos, Himanshu Joshi, Ganesan Venkatasubramanian, John P. John, Neda Jahanshad, Paul M. Thompson

**Affiliations:** Imaging Genetics Center, Mark and Mary Stevens Neuroimaging and Informatics Institute, Keck School of Medicine,University of Southern California, Marina del Rey, CA, United States; Multimodal Brain Image Analysis Laboratory, Translational Psychiatry Laboratory,National Institute of Mental Health and Neuro Sciences (NIMHANS), Bengaluru, Karnataka, India

**Author notes:** Email: jpj.

**Keywords:** tractometry, normative modeling, deep generative model, tractography, diffusion MRI

## Abstract

This study introduces the Deep Normative Tractometry (DNT) framework, that encodes the joint distribution of both macrostructural and microstructural profiles of the brain white matter tracts through a variational autoencoder (VAE). By training on data from healthy controls, DNT learns the normative distribution of tract data, and can delineate along-tract micro- and macro-structural abnormalities. Leveraging a large sample size via generative pre-training, we assess DNT’s generalizability using transfer learning on data from an independent cohort acquired in India. Our findings demonstrate DNT’s capacity to detect widespread diffusivity abnormalities along tracts in mild cognitive impairment and Alzheimer’s disease, aligning closely with results from the Bundle Analytics (BUAN) tractometry pipeline. By incorporating tract geometry information, DNT may be able to distinguish disease-related abnormalities in anisotropy from tract macrostructure, and shows promise in enhancing fine-scale mapping and detection of white matter alterations in neurodegenerative conditions.

## I. Introduction

Diffusion MRI (dMRI) can be used to investigate 3D neural pathways and their microstructural properties in the living human brain, and is collected routinely in large-scale studies of brain aging and neurodegenerative diseases [1]–[3]. Anisotropy and diffusivity measures computed from diffusion tensor imaging (DTI) are often used to characterize WM microstructural properties. Prior work has identified widespread patterns of white matter degeneration in Alzheimer’s disease (AD) using DTI, as reflected by a regionally-specific increase in mean diffusivity and a decrease in fractional anisotropy [4]–[6]. In addition to computing voxel-wise microstructural measures, tractography can be used to delineate long-range WM pathways from the directionality of local diffusion signals, represented by collections of 3D streamlines. Tractometry leverages both DTI and tractography by performing statistics on microstructural scalar measures mapped onto WM tracts. One tractometry method - Bundle Analytics (BUAN) [7] – has been used to identify microstructural abnormalities in AD [8] and bipolar disorder [9], [10] compared to matched controls.

White matter microstructure is widely studied using both voxel-wise and tractometry approaches, but the joint distribution of both microstructural and macrostructural properties is less often studied. In a large multi-site cohort of elder adults, Schilling *et al*. [11] found heterogeneous patterns of age-related macrostructural changes in the brain WM compared to the more homogeneous patterns of microstructural changes. In another study, the same authors reported synchronous micro- and macro-structural changes across the human lifespan [12]. Combining DTI, connectivity and tract shape measures (length, diameter and elongation) derived with a fusion prediction network, Liu *et al*. [13] showed that WM shape measures can improve prediction of cognitive scores in a young healthy cohort. WM micro- and macro-structure have been studied in healthy aging, leading to interest in how these features are independently and jointly altered in neurogenerative conditions such as AD. In addition, given the large number of streamlines and substantial proportion of false positives generated from tractography [14], modeling their joint distribution in a large cohort becomes challenging, but may potentially be tackled using deep learning methods. In our prior work [15], we showed that a convolutional variational autoencoder (ConvVAE) can embed tractography streamlines into a compact latent space and produce new bundles via generative sampling. Autoencoder-based architectures can also be used as a normative model (NM) [16], [17], where they encode statistical distributions of features from a healthy reference population. Deviations from the norm identified by NMs can be used downstream for group difference testing or mapping individual anomalies [18]. At inference time, data from the patient group is passed through the network, and the reconstruction error can be used to conduct group statistics, or for anomaly detection. Using tractography data from the Alzheimer’s Disease Neuroimaging Initiative (ADNI), the ConvVAE-based NM identified 6 WM tracts with along-tract macrostructural anomalies in AD, including the corpus callosum (CC) [19].

In this study, we extend our NM framework and propose *Deep Normative Tractometry (DNT)* to jointly model the 3D geometry of WM tracts and their DTI-derived microstructures. First, we used generative pretraining to encode the complex tract geometric features using a public multi-site dMRI dataset of healthy subjects from Europe and North America. To test the model’s ability to generalize to a new population, we applied transfer learning to adapt the model to data from an independent cohort acquired in India. We report along-tract microstructure and macrostructure deviations computed from DNT in mild cognitive impairment (MCI) and AD. To further investigate the effect of incorporating WM geometry, we compare microstructural findings between the BUAN tractometry pipeline [7] and DNT.

## II Data

We analyzed 3T diffusion MRI (dMRI) data in a sample of 302 participants from two scanners from the NIMHANS (National Institute of Mental Health and Neuro Sciences) cohort (see Table I) in India. The dataset consists of neuroimaging data from 123 cognitively normal participants (CN), 89 diagnosed with mild cognitive impairment (MCI) and 90 with AD. Diffusion-weighted image (DWI) acquisition for the Philips 3T Ingenia scanner was performed using a single-shot, DWI echo-planar imaging sequence (TR=7441 ms, TE=85 ms,TA=630 s, voxel size: 2 *×* 2 *×*2mm^3^, 64 slices, flip angle=90^*°*^, FOV 224mm) [20]. DWI acquisition for the Siemens 3T Skyra was performed using a single-shot, DWI echo-planar imaging sequence (TR=8400 ms, TE=91 ms, TA=546 s, FOV 240mm) [21], [22]. For both scanners, transverse sections of 2-mm thickness were acquired parallel to the anterior commissure-posterior commissure (AC-PC) line, and diffusion weighting was encoded along 64 independent orientations, using *b*-value of 1000s/mm^2^. Preprocessing steps for all DWI volumes included: denoising using local principal component analysis [23], Gibbs ringing removal [24], [25], correction of susceptibility-induced distortion [26], eddy currents [27] and bias field inhomogeneity [28]. Diffusion tensors were fitted with non-linear least-squares to produce fractional anisotropy (FA) and medial diffusivity (MD) scalar maps in the DWI native space. Fiber orientations were reconstructed using Robust and Unbiased Model-BAsed Spherical Deconvolution (RUMBA-SD) [29], [30]. Particle filtering tracking [31] was using to generate whole-brain tractograms (WBT), with 8 seeds per voxel generated from the WM mask, step size of 0.2 mm, angular threshold of 30°, and the continuous map stopping criterion.

**TABLE I.**
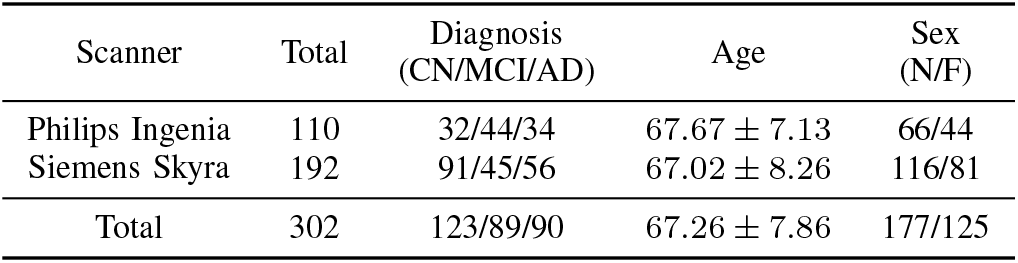
NIMHANS dataset description, by scanner.

| Scanner | Total | Diagnosis<br>(CN/MCI/AD) | Age | Sex<br>(N/F) |
| --- | --- | --- | --- | --- |
| Philips Ingenia | 110 | 32/44/34 | $67.67 \pm 7.13$ | 66/44 |
| Siemens Skyra | 192 | 91/45/56 | $67.02 \pm 8.26$ | 116/81 |
| Total | 302 | 123/89/90 | $67.26 \pm 7.86$ | 177/125 |

The pre-training dataset was composed of 198 single-shell dMRI volumes from the publicly available TractoInferno training dataset [32], acquired on 3T scanners from 6 sites. Constrained spherical deconvolution [33] and deterministic tracking [34] were used to generate WBTs. For both the datasets, DIPY’s [34] auto-calibrated RecoBundles [7], [35] was used to segment thirty WM bundles in the native and MNI (Montreal Neurological Institute) space [36].

## III Method

### A. BUndle ANalytics (BUAN)

The BUAN tractometry pipeline was used to quantify along-tract microstructural differences in MCI and AD from the NIMHANS cohort (see Figure 1). For each subject, FA and MD scalar maps computed in the DWI native space were mapped to all streamline vertices in 30 bundles in the same space. Bundles extracted in the MNI space were used to create 100 along-tract segments, aligned across subjects. For each group comparison (MCI vs. CN and AD vs. CN), we computed group statistics using a linear mixed model (LMM), where diagnosis, age, and sex were modeled as fixed effects, and subject and scanner were modeled as random effects. We then corrected for multiple comparisons using the false discovery rate (FDR) method [37] for 100 segments on each bundle; we report *p*_FDR_ *<* 0.05.

**Fig. 1.**
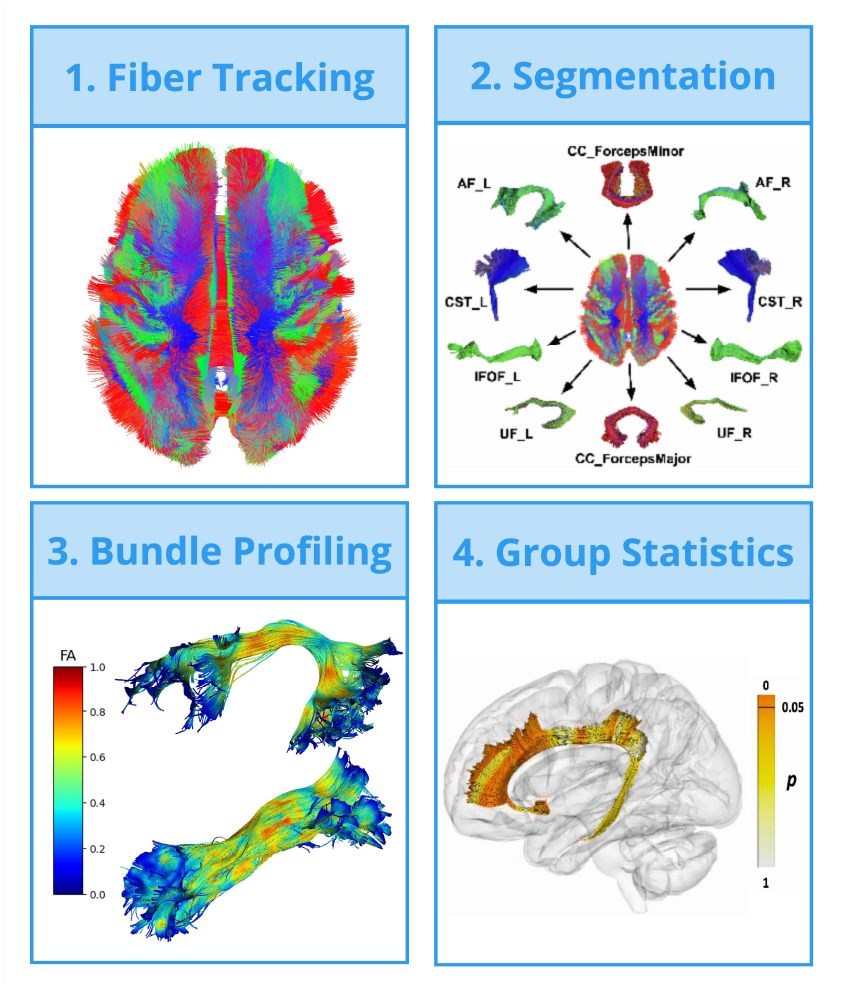
The BUAN tractometry pipeline: 1. Whole-brain tractograms are generated from dMRI; 2. WM bundles are segmented from whole-brain tractograms; 3. Bundle profiles are created by projecting microstructural measures onto segmented tracts, and linear mixed models are applied to detect group differences; 4. significant group differences are mapped onto the bundle for visualization.

### B. Deep Normative Tractometry (DNT)

The DNT pipeline consists of two training stages on healthy controls to create the normative model, followed by model inference, along-tract anomaly mapping, and statistics (see Figure 2). Each streamline from 30 bundles in the MNI space (from both the TractoInferno and NIMHANS dataset) was first resampled to 128 equidistant points; DTI-derived FA and MD scalars were then mapped to each point on the corresponding native space bundle to create the bundle profile. Combined with the 3D spatial coordinates, each streamline,of size 128 *×* 5, was used as input to the ConvVAE model to jointly model both bundle geometry and microstructure.

**Fig. 2.**
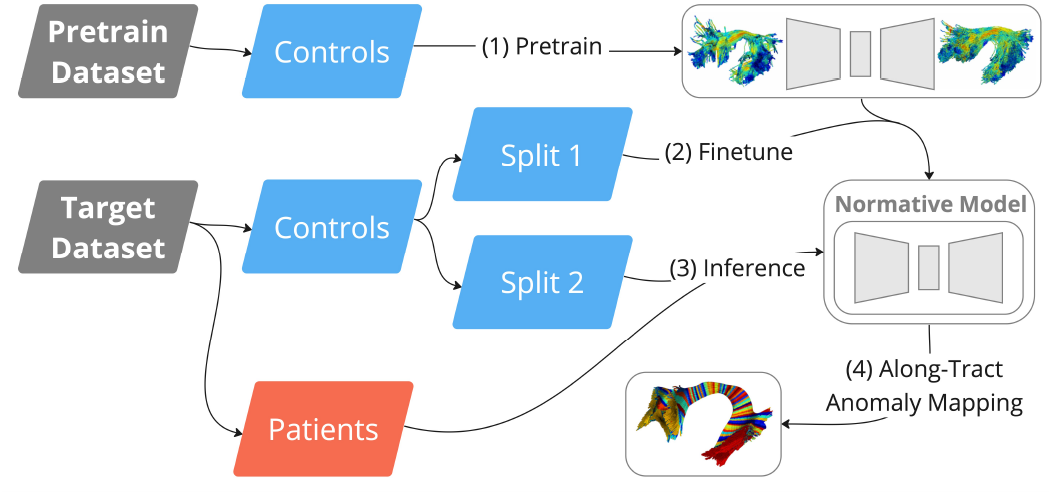
The Deep Normative Tractometry (DNT) framework.

The model had an asymmetric architecture, with a 4-layer encoder, a 3-layer decoder, and a latent dimension of 64. Depthwise separable convolution [38] was used in each layer to accommodate a deeper feature space with efficient use of model parameters. The model, containing 454,742 trainable parameters, was pretrained for 29,000 steps, with a batch size of 512 on the TractoInferno dataset. Fifty CN subjects from the NIMHANS dataset, randomly selected and stratified by sex and scanner, were then used to fine-tune the model for 5000 steps. At inference time, bundle profiles from the remaining CN, MCI, and AD subjects were passed through the model to obtain their point-by-point reconstruction. Each bundle and its reconstruction were then divided into 100 along-tract segments using the same approach described in Section III-A. The anomaly scores are computed using the Mean Absolute Error (MAE) for each segment, and each feature - 3D coordinates (shape), FA, and MD. We then corrected for age, sex, and scanner effect using linear regression. With the assumption that the anomaly score is higher in the MCI and AD group for all features, we conducted MCI vs. CN and AD vs. CN group statistics on the anomaly scores using a 1-tailed Mann-Whitney *U* test. We corrected for multiple comparisons using the same FDR approach as the one used for BUAN.

## IV Results

To compare results for both BUAN and DNT, we plot the number of along-tract segments that showed significant group differences after FDR correction for each bundle, feature (shape, FA and MD) and diagnosis (MCI, AD) in Figure 3. Most notably, we see widespread MD differences between AD and CN - identified by both methods - in the commissural, association, and projection tracts, followed by moderate alignment in MD differences between MCI and CN. In bundle segments that show significant differences, BUAN reports higher MD and DNT also reports higher MD anomaly scores in both MCI and AD groups when compared to CN. In Figure 4, we plot the along-tract − log_10_(*p*_FDR_) values from AD vs. CN comparisons in the commissural tracts (CC mid, *forceps major* and *forceps minor*). DNT reveals MD abnormality patterns similar to BUAN along the length of the tracts, even when accounting for shape changes.

**Fig. 3.**
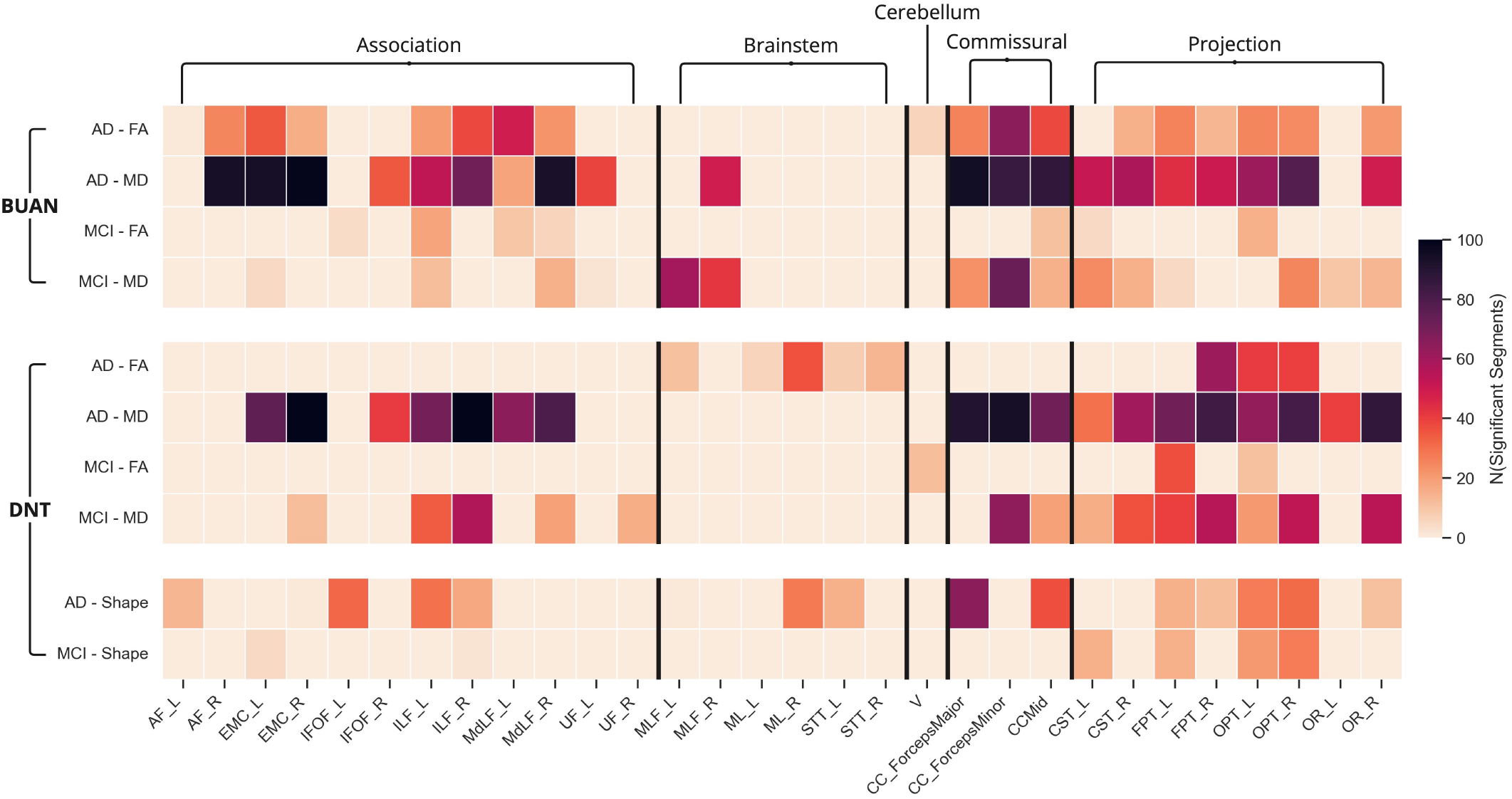
The number of along-tract segments out of 100 that showed significant group differences after FDR correction for both BUAN and DNT. The tracts description can be found in [36].

**Fig. 4.**
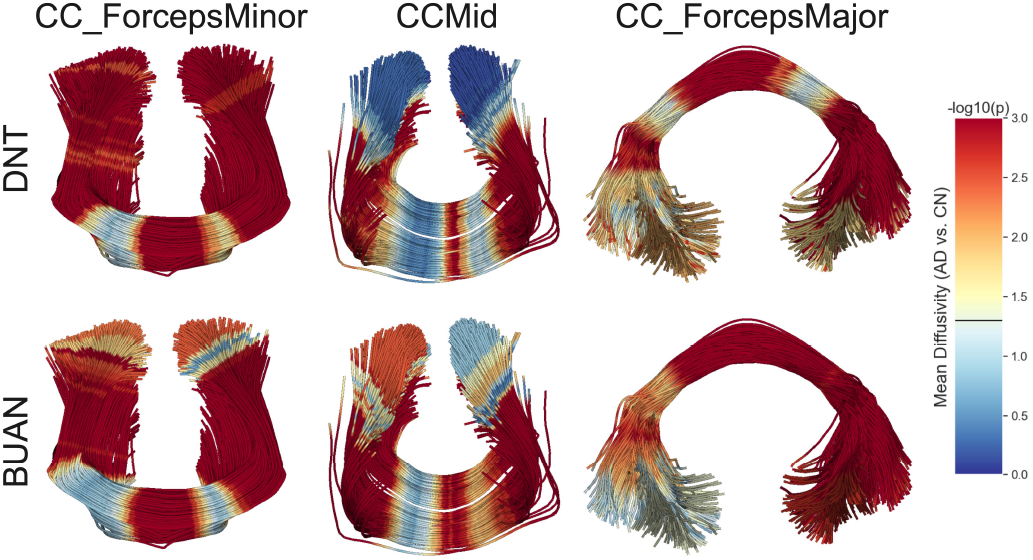
Along-tract − log_10_(*p*_FDR_) from AD vs. CN comparison of MD in the commissural tracts from both DNT and BUAN. The threshold for statistical significance of *p*_FDR_ is plotted as a horizontal line on the colorbar.

BUAN reveals more along-tract regions with significant FA differences in both MCI and AD than DNT, with few alignments between the two methods. However, by incorporating information on 3D spatial coordinates (i.e., the tract’s overall geometrical shape), DNT identifies a moderate number of tracts with significant shape differences, primarily in the projection and commissural tracts. Here, we show examples of the *forceps major* (CC ForcepsMajor) and the left occipitopontal tract (OPT L) in Figure 5. In the AD vs. CN comparison, BUAN identifies significant FA differences in both bundles, whereas DNT identifies significant group differences of FA anomalies only in OPT L, but shape anomalies in both. Patterns of FA group differences in OPT L from both methods are similar to that of the shape shown from DNT. In the *forceps major*, patterns of FA differences from BUAN are aligned with the profile of group differences in shape anomalies from DNT, but we observe no difference in FA when the macrostructure is jointly modeled with the microstructure. Accounting for shape changes, FA differences in OPT L in AD subjects may reflect aspects of the underlying loss of fiber integrity in AD, and FA differences in the *forceps major* observed in BUAN may be attributed to disease-related macrostructural changes. Prior work using structural MRI and fixel-based analysis identified structural atrophy of the corpus callosum in people with AD, possibly associated with ventricular dilation [39], [40]. Due to the partial volume effect, particularly in the CC tracts [41], disease-related macrostructural changes could contribute to changes in FA when the underlying fiber geometry is not accounted for.

**Fig. 5.**
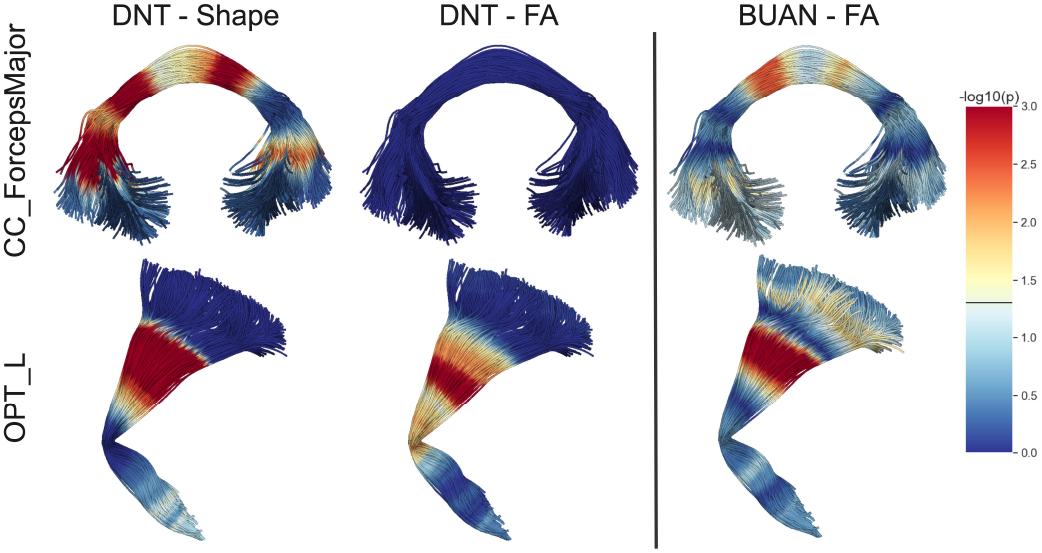
Along-tract − log_10_*major*(*p*_FDR_) from AD vs. CN comparison in the *forceps*(CC ForcepsMajor) and the left occipito-pontine tract (OPT L). Themeasures shown here are shape differences reported by DNT, and FA differences reported by both DNT and BUAN.The threshold for statistical significance of *p*_FDR_ is plotted as a horizontal line on the colorbar.

We further highlight shape and MD differences detected by DNT in both MCI and AD compared to CN in Figure 6. The along-tract − log_10_(*p*_FDR_) values and Cohen’s *d* effect sizes are plotted for 2 associative and 2 projections WM tracts. Both macrostructural and diffusivity changes are more widespread along tracts in AD - with a medium to large effect compared to MCI, where segments with significant differences in MCI are also identified in AD. Patterns of shape differences are more localized along-tract, whereas patterns of MD differences are also more widespread. MD results reported by DNT are also consistent with those previously reported by BUAN applied to an MCI cohort from ADNI3 [2], [8].

**Fig. 6.**
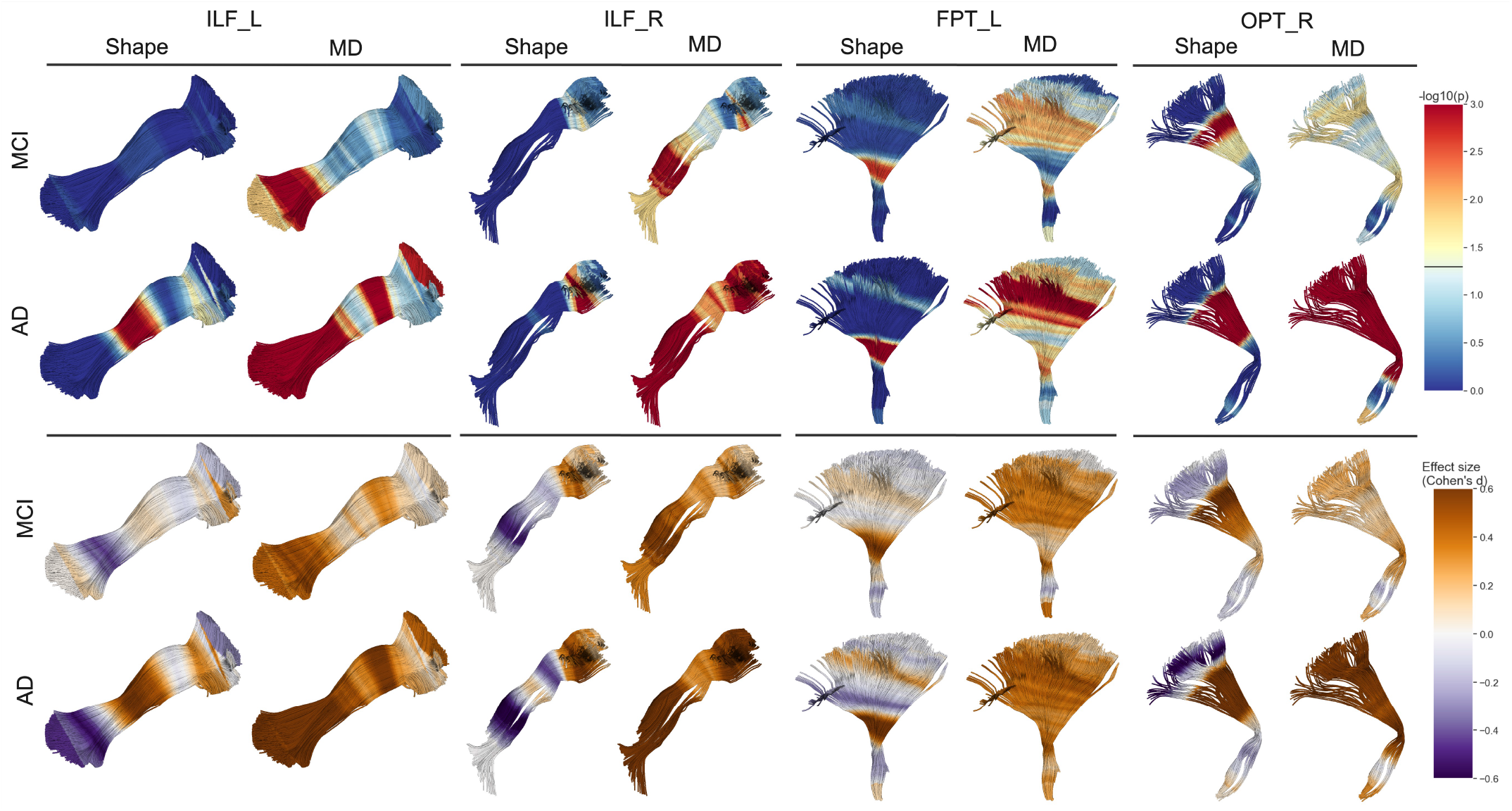
Along-tract shape and MD group differences identified by DNT in MCI, AD versus CN subjects. The four tracts shown are the left and right inferior longitudinal fasciculus (ILF), the left fronto-pontine tract (FPT), and the right occipito-pontine tract (OPT)

## V. Discussion

In this study, the DNT framework is compared with the BUAN tractometry pipeline for detecting microstructural anomalies in MCI and AD subjects. We found that effects on MD showed widespread alignment across both methods compared to FA in both the MCI and AD group. One surprising finding is that when accounting for bundle macrostructure, DNT detects no difference in FA in the commissural tracts. This could suggest that changes in FA may be confounded by the underlying tract structural abnormalities, and our framework could help to alleviate the partial volume effect by integrating non-local information from streamlines. In the current DNT framework, reconstruction error is used as a proxy for shape anomalies, and it does not provide a detailed characterization of structural changes, such as fiber length, curvature, cross-section, and density. Future work will investigate advanced shape metrics to quantify multi-scale shape differences [42].

As crossing fibers are widespread in the brain [43], [44], both micro- and macro-structural abnormalities detected by DNT in one WM tract could still be due, partly or fully, to differences in another fiber population, even when accounting for bundle geometry. Due to the underlying limitations of DTI and tractography [45], it remains challenging to disentangle the effect of multiple fiber populations in a single voxel or fixel [44] from diseased-related bundle macrostructural and microstructural changes with high specificity.

Aside from directly modeling bundle geometry, DNT is pretrained on data that BUAN has not seen, and could contribute to the differences in signals detected by both methods. The pretraining step helps the model to learn the complex distributions of WM macrostructure and microstructure, so this may prevent overfitting and produce better characterizations of the target cohort after fine-tuning, especially when data is limited in the target cohort. TractoInferno was selected as the pretraining dataset in this study, as it contains multi-site single-shell acquisitions and has been carefully quality controlled. Intuitively, knowledge transfer may be more effective when the source and target data distributions are more similar. However, in the context of tractography, including data from high angular resolution diffusion imaging (HARDI) acquisitions, such as the Human Connectome Project (HCP) [46], in the pre-training dataset may help the model better distinguish complex fiber populations. Other single-shell metrics beyond DTI and more advance multi-shell metrics may also provide a richer feature set. Future work will include more advanced diffusion models such as the tensor distribution function (TDF) [47], neurite orientation dispersion and density imaging (NODDI) [48] and diffusion kurtosis imaging (DKI) [49], which may be able to evaluate AD-related abnormalities with greater sensitivity and specificity [6], [50], [51]. In future, we will also investigate the effect of pre-training, data sources, feature sets, sample size, and fine-tuning techniques.

The normative tractometry framework in DNT can better account for variability within the group, as opposed to testing for mean differences in the traditional case-control schema. Generative models, which explicitly model the marginal or the joint distribution - as opposed to discriminative models - are well-suited for normative modeling. In DNT, the VAE model learns the joint distributions of shape, MD, and FA from CN subjects at the training stage. To compare with the LMM results from BUAN, we used the along-tract anomaly scores from DNT for group difference testing. However, a normative framework such as DNT can also be applied for single subject anomaly detection, or regression analysis with non-imaging phenotypes [18].

## VI Conclusion

In this study, we proposed the DNT framework to jointly model the 3D geometry and microstructural profiles of WM tracts using a deep generative model based on a convolutional VAE. The model learns the normative distribution by training on data from healthy controls, and can be used to map along tract micro- and macro-structural abnormalities. DNT leverages a large sample size using generative pretraining, and we tested its generalizability using transfer learning on an independent cohort. DNT was able to identify widespread along-tract diffusivity abnormalities in MCI and AD, with high levels of agreement with the state-of-the-art BUAN tractometry pipeline. By accounting for the 3D tract geometry, DNT may be able to disentangle disease-related group differences in anisotropy from macrostructural abnormalities.

## VII Acknowledgments

This research was supported by the NIH (National Institutes of Health) under the AI4AD project grant U01 AG068057, grant numbers P41 EB015922, and RF1 AG057892. We would like to acknowledge funding support from the NIH grant R01 AG060610 and the Department of Science and Technology, Govt. of India, grant nos. DST-SR/CSI/73/2011 (G); DST-SR/CSI/70/2011 (G); and DST/CSRI/2017/249 (G).

